# NanoSim: nanopore sequence read simulator based on statistical characterization

**DOI:** 10.1101/044545

**Authors:** Chen Yang, Justin Chu, René L Warren, Inanç Birol

**Affiliations:** Faculty of Science, University of British Columbia, Vancouver, Canada; Genome Science Centre, British Columbia Cancer Agency, Vancouver, Canada; Department of Medical Genetics, University of British Columbia, Vancouver, Canada; School of Computing Science, Simon Fraser University, Burnaby, Canada.

**Author notes:** **Availability:** NanoSim is written in Python and R. The source files and manual are available at the Genome Sciences Centre website. www.bcgsc.ca/platform/bioinfo/software/nanosim.

## Abstract

**Motivation:** In 2014, Oxford Nanopore Technologies (ONT) announced a new sequencing platform called MinION. The particular features of MinION reads – longer read lengths and single-molecule sequencing in particular – show potential for genome characterization. As of yet, the pre-commercial technology is exclusively available through early-access, and only a few datasets are publically available for testing. Further, no software exists that simulates MinION platform reads with genuine ONT characteristics.

**Results:** In this article, we introduce NanoSim, a fast and scalable read simulator that captures the technology-specific features of ONT data, and allows for adjustments upon improvement of nanopore sequencing technology.

## 1 Introduction

DNA sequencing is dominated by sequencing-by-synthesis technologies, and mature second generation systems such as those from Illumina Inc. are amongst the most widely adopted. In recent years, third generation single molecule sequencing using nanopore-based technologies have emerged, with promises of longer reads and lower cost. Launched by Oxford Nanopore Technologies (ONT) in April 2014, the MinION sequencer, which is currently in pre-release testing, stands out among existing third generation sequencing technologies due to its ability to generate ultra-long reads, albeit with high error rates. For example, the *S. cerevisiae* dataset has an average read length of 5,473 bp, and maximum reaching 147 kbp, whereas the sequence identity is 64% for 1D reads and 75% for 2D reads (Goodwin *et al*., 2015), 1D and 2D referring to interrogation of a DNA molecule template once or twice, respectively.

Long nanopore reads hold great potential for *de novo* assembly and transcriptome analysis as they can span more repetitive regions and multiple exon junctions, or even entire transcripts. However, the error-prone reads pose new challenges to algorithm design (Jain *et al*., 2015). As it is the case for other sequencing platforms (Hu *et* al., 2012), a read simulator designed specifically for ONT reads is desirable in order to develop and benchmark new algorithms, with the aim to harness the full potential of this new sequencing platform. Currently, however, no state-of-the art DNA sequence simulator emulates the properties of ONT reads.

Here, we introduce NanoSim, a nanopore sequence read analysis and simulation pipeline. The tool analyzes ONT reads from experimental data to model read features, such as error profiles and length distributions, and uses these features to generate *in silico* reads for an input reference. We show that the statistical models NanoSim uses remain valid as the nanopore sequencing technology evolves.

## 2 Methods

NanoSim is implemented using R for error model fitting, and Python for read length analysis and simulation (Supplementary Fig. S1). The first step of NanoSim is read characterization, which provides a comprehensive alignment-based analysis, and generates a set of read profiles serving as the input to the next step, the simulation stage. The simulation tool uses the model built in the previous step to produce *in silico* reads for a given reference genome. It also outputs a list of introduced errors, consisting of the position on each read, error type and reference bases.

The modeling stage of NanoSim takes a reference and a training read set in FASTA format as input. The reads are aligned to the reference genome using LAST with tuned parameters (‘-r 1-q 1-a 1-b 1’) (Frith *et al.*, 2010). Alternatively, the tool also allows the input of an alignment file in MAF format. If not unique, the best alignment of each read is chosen based on alignment length to avoid the influence of mis-alignments due to repeat regions.

Based on alignment results, training reads are classified into two types, aligned reads and unaligned reads. For aligned reads, typically only a middle region can be aligned, leaving the flanking head and tail regions soft-clipped from alignments. The length distribution of head and tail regions exhibits a multimodal pattern. The full read length distribution can be characterized by two empirical distributions: one for the length of the aligned regions, the second for the ratio of alignment lengths to read lengths. Length distributions of unaligned reads are also generated to simulate unaligned reads. The perfect flag of NanoSim can generate perfect reads with no errors, relying on the full-length distribution of aligned reads.

Sequencing errors on the aligned region share similar patterns among different datasets, which can be described by statistical mixture models (Warren *et al.*, 2015):

Mismatch: *P*_*m*_ ∼ α_*m*_ Poisson (*λ*_*m*_) + (1 - α_*m*_) Geometric (*p*_*m*_)

Insertion: *P*_*i*_ ∼ α_*i*_ Weibull (λ_*i*_, *κ*_*i*_) + (1 - α_*i*_) Geometric (*p*_*i*_)

Deletion: *P*_*d*_ ∼ α_*d*_ Weibull (λ_*d*_, *κ*_*d*_) + (1 - α_*d*_) Geometric (*p*_*d*_)

Here α_*m/i/d*_ Î (0, 1) are mixture parameters, *p*_*m/i/d*_ are the event probabilities in the geometric distributions, λ_*m*_ is the expected value of the Poisson distribution, and λ_*i/d*_ and *κ*_*i/d*_, respectively, are the scale and shape parameters of the Weibull distributions.

The mixture model describes stretches of substitution errors as being distributed according to Poisson distribution, whereas indels following Weibull distributions. All error modes have a second component of geometric distribution, which we postulate describes stochastic noise. Moreover, the structures of these models remain unchanged when we use an alternative aligner (BWA-MEM with the ‘-x ont2d’ option) (Li *et al*., 2013).

The model parameters and error profiles for the tested datasets are provided with the software download package, and can be directly used for simulation.

During simulation, the lengths of errors are drawn from the statistical models, and the error types are determined by a Markov chain, simulating the transitional probability between two consecutive errors (Supplementary Fig. S2). Interval lengths between errors are observed to be auto-correlated, and justifies the use of a Markov chain to model interarrival times between errors (Supplementary Fig. S3).

Reads that are unaligned are more difficult to characterize. Rather than assuming them to be random sequences, we extract sequences from the reference, but use an arbitrarily high error rate compared to the aligned reads. We pick the length of each error in these reads from the same mixture models as the aligned reads, and randomly place them on the simulated sequence.

Another feature of NanoSim is that it is able to simulate either circular or linear genomes. A read extracted from a circular genome can start from any position and may wrap around. If the length of a read is longer than the length of the whole genome, which is unlikely but possible for a plasmid or viral genome, it will be truncated to the genome length. For a linear genome to maintain a read length distribution similar to the training profile, NanoSim will only extract reads from chromosomes that are longer than the read length.

The k-mer bias, especially the deficiency of long homopolymers, has been well-studied (Loman *et al*., 2015). As a DNA molecule with a stretch of homopolymer sequence traverses through a nanopore, the change in electric current is not detectable or fails to be interpreted by the base-calling algorithm, leading to a deficient representation of homopolymers longer than the number of bases that can fit in the nanopores. The k-mer bias mode of NanoSim compresses all homopolymers longer than *n* into *n*-mers (default *n*=5), simulating the process of base-calling. The under or over representation of other k-mers is not supported in the current version of NanoSim. However, we expect this sequencing bias to be addressed by the vendor in the future, given the improvement of the R7.3 chemistry compared to the previous R7 chemistry (Supplementary Fig. S4).

Using an *E. coli* dataset, it has been reported that the GC content of 2D reads is very close to the reference, and that this has a minor effect on sequencing error rates (Karlsson *et al*., 2015). In prior work, we have also observed that substitution errors are not uniform, with a weak bias towards G and C (Warren *et al*., 2015). Since the underlying mechanism causing this bias is unclear, this pattern is not reflected in the NanoSim synthetic reads.

## 3 Results and discussion

Four datasets were used to benchmark NanoSim, including three *E. coli* datasets and one *S. cerevisiae* dataset (Table 1). Generally, 2D reads have higher quality than 1D reads, and are more frequently used in downstream analyses. As such, we only tested NanoSim on 2D reads. All tests were performed on a single machine with 8-core Intel i7-4770 CPUs @ 3.40GHz and 8 GB total RAM.

**Table. 1.**
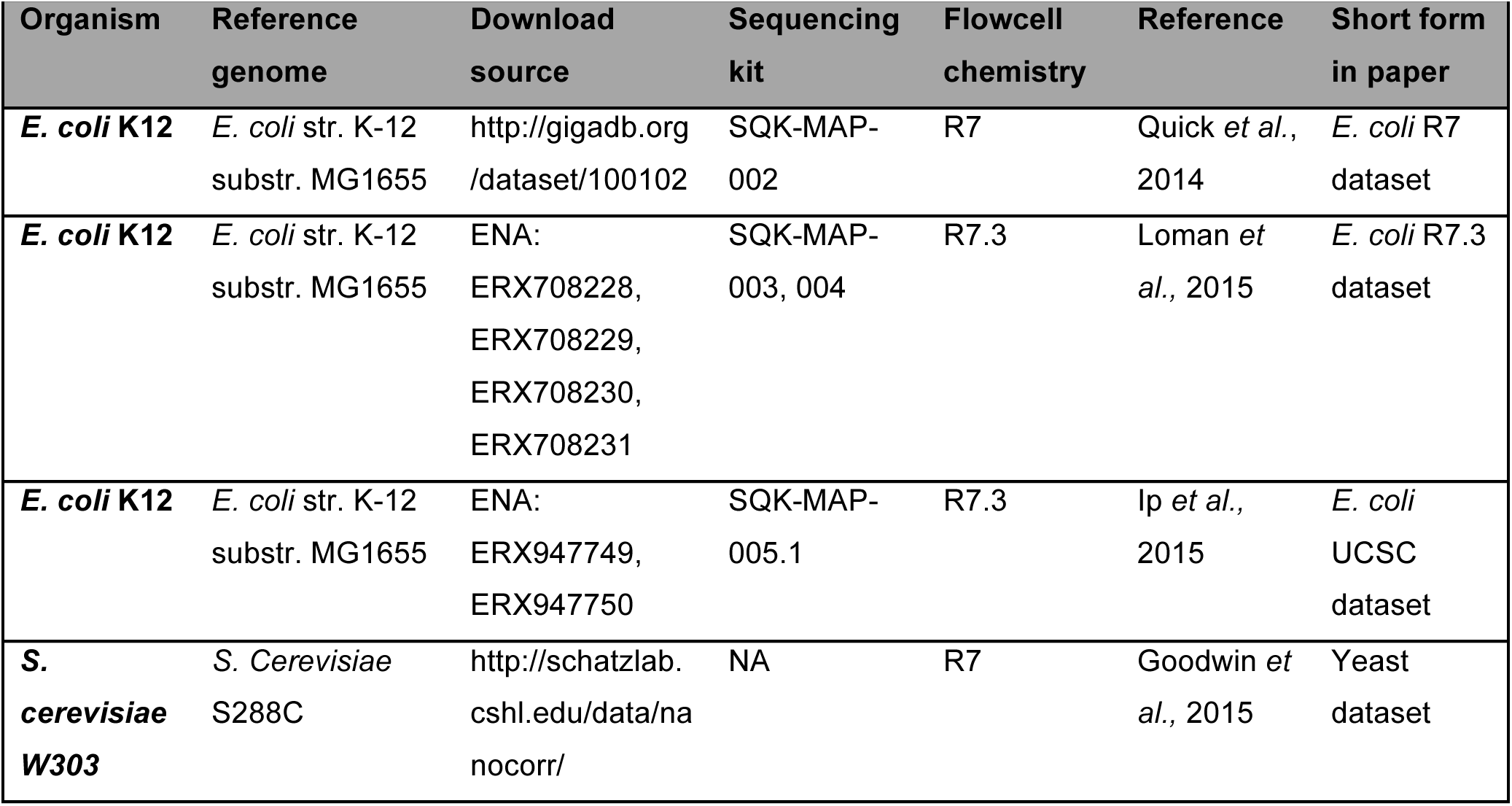
Datasets used for benchmarking

| Organism | Reference genome | Download source | Sequencing kit | Flowcell chemistry | Reference | Short form in paper |
| --- | --- | --- | --- | --- | --- | --- |
| ***E. coli* K12** | *E. coli* str. K-12 substr. MG1655 | http://gigadb.org/dataset/100102 | SQK-MAP-002 | R7 | Quick *et al.*, 2014 | *E. coli* R7 dataset |
| ***E. coli* K12** | *E. coli* str. K-12 substr. MG1655 | ENA: ERX708228, ERX708229, ERX708230, ERX708231 | SQK-MAP-003, 004 | R7.3 | Loman *et al.*, 2015 | *E. coli* R7.3 dataset |
| ***E. coli* K12** | *E. coli* str. K-12 substr. MG1655 | ENA: ERX947749, ERX947750 | SQK-MAP-005.1 | R7.3 | Ip *et al.*, 2015 | *E. coli* UCSC dataset |
| ***S. cerevisiae* W303** | *S. Cerevisiae* S288C | http://schatzlab.cshl.edu/data/nanocorr/ | NA | R7 | Goodwin *et al.*, 2015 | Yeast dataset |

### 3.1 Speed and memory

The runtime of NanoSim scales up linearly with the number of reads (Supplementary Fig. S5), and the memory requirement depends on the length of the reference sequence. For example, the *E. coli* UCSC dataset contains 45,049 2D pass reads with an average length of 7,067 bp. Excluding read alignments, the characterization stage of NanoSim took 22m:32s, and the peak memory usage was 2.68 GB. Simulating 20,000 *E. coli* reads took 4m:39s; peak memory usage was 120 MB.

### 3.2 Simulation result and tool comparison

The error models derived from the characterization stage are consistent across both chemistries and organisms (Supplementary Tables S1-S3). Assessing the goodness of fit via a Kolmogorov–Smirnov test, we observed that base call error distributions were statistically identical to their fitted models with p-value > 0.05 (Supplementary method).

Currently, there are simulators that could potentially simulate Nanopore-like reads, such as PBSIM (Ono et al., 2013), ReadSim (Lee *et al*., 2014) and FASTQSim (Shcherbina, 2014). Among these, PBSIM is designed to simulate reads from Pacific Biosciences (PacBio) sequencers, which also produce long, yet error-rich reads. FASTQSim is a platform-independent simulator that can theoretically simulate any NGS datasets. ReadSim 1.6 is the only simulator, which advertises the ability to simulate ONT reads.

Thus to evaluate the accuracy of NanoSim, we conducted comparisons only with ReadSim. In each experiment on the four datasets in Table 1, 20,000 synthetic reads were generated by NanoSim and ReadSim. Since ReadSim is not capable of simulating genomes with multiple chromosomes, for the yeast dataset we linked the yeast chromosomes with a single “N” in between before simulation, and discarded synthetic reads containing “Ns”. Simulated reads were aligned back to the reference genome and analyzed using the characterization tool of NanoSim.

ReadSim simulates read lengths through a sample-based method or a Gaussian-model-based method. The sample-based method was used here and fed with the empirical lengths of all reads regardless of alignment results. After simulation, over 99.9% synthetic reads produced by ReadSim can be aligned to the reference, while raw ONT datasets and NanoSim reads agree on the alignment rates ranging from 82.83% to 99.68% for these four datasets.

The length of consecutive perfect/error bases of simulated reads were plotted together along with their raw experimental read counterparts (Fig. 1A, Supplementary Fig. S6A, S7A, S8A). We observed that the ReadSim reads deviate further away from experimental data because they were simulated with uniformly distributed errors and randomly chosen error length.

**Fig. 1.**
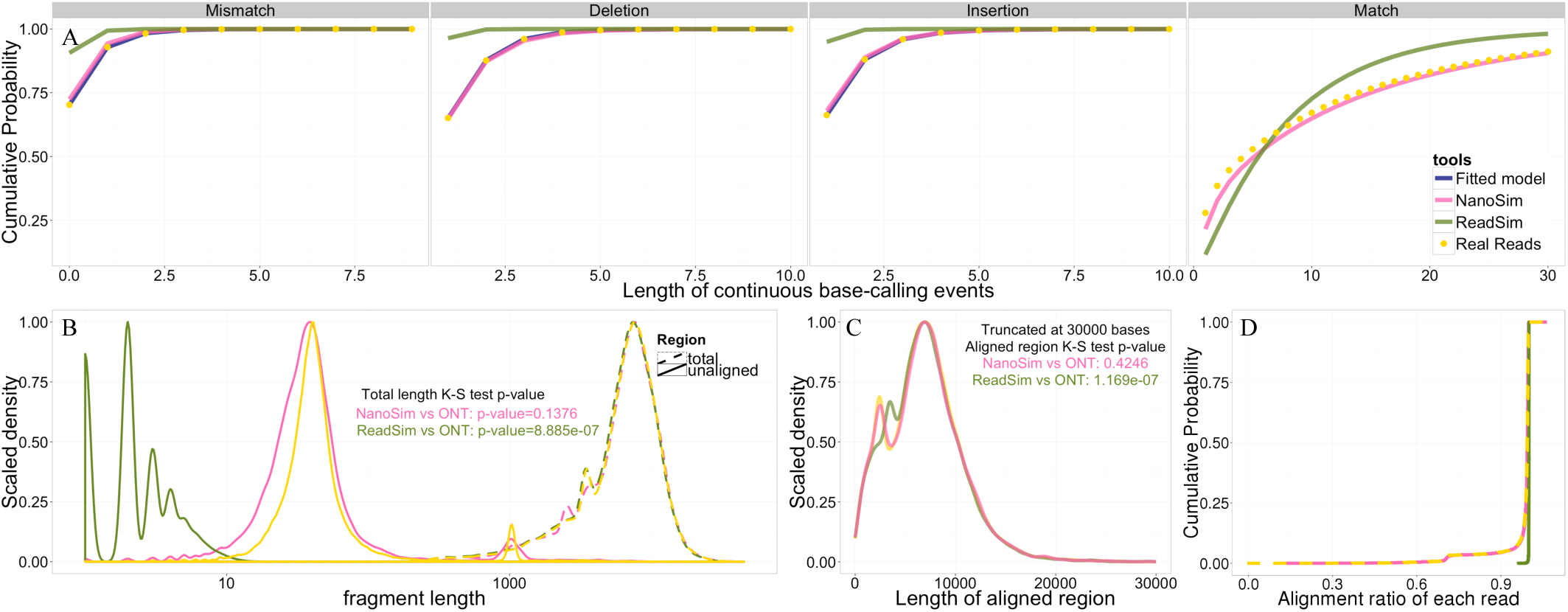
NanoSim and ReadSim simulation results compared with UCSC E. coli experimental reads. (A) The four plots on the upper panel are cumulative density plots of error match events and error events. (B) Length density plot of unaligned regions and total read lengths of aligned reads. (C) Length density plot of aligned regions on each read. (D) Cumulative density plot of the alignment ratio of each read.

Statistically speaking, for all aligned reads, the lengths of the whole read and aligned regions of NanoSim reads and ONT reads are drawn from the same distributions (Fig. 1B, 1C, Supplementary Fig. S6B, S6C, S7B, S7C, S8B, S8C). The distribution of aligned regions also exhibits bi-modal pattern with two peaks. Whereas, the only length distribution ReadSim re-produces well is the full length distribution of aligned reads on *E. coli* R7.3 dataset (Supplementary Fig. S7B).

Since the lengths of ReadSim reads are drawn from the empirical data points directly, and over 99.9% ReadSim reads can be aligned, the full-length distribution of aligned ReadSim reads is identical to the full-length distribution of all ONT reads. By comparing the total length density of ONT and ReadSim aligned reads, we observe that the length distributions of aligned reads and unaligned reads do not agree on all datasets except for *E. coli* R7.3 (Supplementary Fig. S7B).

The lengths of unaligned regions are determined by the alignment ratio of each read. NanoSim performed better on *E. coli* R7 and yeast datasets than *E. coli* R7.3 and *E. coli* UCSC datasets, generating almost identical distributions of alignment ratio as the raw ONT reads (Fig. 1D, Supplementary Fig. S6D, S7D, S8D). This leads to similar statistical test results on the distribution of unaligned head and tail regions (Fig. 1B, Supplementary Fig. S6B, S7B, S8B). The unaligned regions on experimental ONT reads also have two peaks, and for *E. coli* UCSC dataset, they centered at 40 bp and 1000 bp (Fig. 1B). NanoSim reads overlap with these two peaks on all four datasets, whereas ReadSim reads have much shorter unaligned regions. The head and tail regions are not profiled and thus not recovered by ReadSim.

## 4 Conclusions

In summary, NanoSim mimics ONT reads well, true to the major features of the emerging ONT sequencing platform, in terms of read length and error modes. The independent profiling module grants users the freedom to characterize their own ONT datasets, which is expected to perform consistently upon the improvement of nanopore sequencing technology, as the shapes of the error models hold among different datasets. NanoSim will immediately benefit the development of scalable NGS technologies for the long nanopore reads, including genome assembly, mutation detection, and even metagenomic analysis software. Currently, no human genome-size data sequenced by nanopore technologies are yet available. With the help of NanoSim, bioinformatics software developers can easily test the scalability of their tools using simulated reads. Moreover, a mixture of *in silico* genomes simulating a microbiome will be helpful for benchmarking algorithms with application in metagenomics, including functional gene prediction, species detection, comparative metagenomics, clinical diagnosis. As such, we expect NanoSim to have an enabling role in the field.

## Acknowledgements

We thank Genome Canada, Genome British Columbia, British Columbia Cancer Foundation, and University of British Columbia for their financial support. The work is also partially funded by the National Institutes of Health under Award Number R01HG007182. The content of this work is solely the responsibility of the authors, and does not necessarily represent the official views of the National Institutes of Health or other funding organizations.

*Conflict of Interest:* none declared.

## References

Frith, M. C. et al. (2010). Parameters for accurate genome alignment. BMC bioinformatics, 11(1), 80.

Goodwin, S. et al. (2015). Oxford Nanopore sequencing and de novo assembly of a eukaryotic genome. Genome Res. 2015 Nov;25(11):1750–6.

Hu, X. et al. (2012). pIRS: Profile-based Illumina pair-end reads simulator. Bioinformatics, 28(11), 1533–1535.

Ip, C. L. et al. (2015). MinION Analysis and Reference Consortium: Phase 1 data release and analysis F1000Res. 2015 Oct 15;4:1075.

Jain, M. et al. (2015). Improved data analysis for the MinION nanopore sequencer. Nature methods, 12(4), 351.

Karlsson, E. et al. (2015). Scaffolding of a bacterial genome using MinION nanopore sequencing. Scientific reports, 5.

Lee, H. et al. (2014). Error correction and assembly complexity of single molecule sequencing reads. bioRxiv, 006395.

Li, H. (2013). Aligning sequence reads, clone sequences and assembly contigs with BWA-MEM. arXiv preprint arXiv:1303.3997.

Loman, N. J., et al. (2015). A complete bacterial genome assembled *de novo* using only nanopore sequencing data. Nature Methods 12, 733–735

Ono, Y. et al. (2013). PBSIM: PacBio reads simulator—toward accurate genome assembly. Bioinformatics, 29(1), 119–121.

Quick, J. et al. (2014). A reference bacterial genome dataset generated on the MinION^™^ portable single-molecule nanopore sequencer. GigaScience, 3(1), 22.

Shcherbina, A. (2014). FASTQSim: platform-independent data characterization and in silico read generation for NGS datasets. BMC research notes, 7(1), 533.

Warren, R. L. et al. (2015). LINKS: Scalable, alignment-free scaffolding of draft genomes with long reads. GigaScience, 4(1), 1–11.

